# Individual Differences in Belief Updating and Phasic Arousal Are Related to Psychosis Proneness

**DOI:** 10.1101/2024.01.14.575567

**Authors:** Peter R Murphy, Katarina Krkovic, Gina Monov, Natalia Kudlek, Tania Lincoln, Tobias H Donner

## Abstract

Many decisions entail the updating of beliefs about the state of the environment, a process that may go awry in psychosis. When environments are subject to hidden changes in their state, optimal belief updating requires non-linear modulation of sensory evidence, which may be subserved by pupil-linked, phasic arousal. Here, we analyzed behavior and pupil responses during evidence accumulation in a changing environment in a community sample of human participants and assessed their subclinical psychotic experiences (psychosis proneness). Subjects most prone to psychosis showed overall less flexible belief updating profiles, with diminished weighting of late evidence. These same subjects also exhibited overall smaller pupil responses and less reliable pupil encoding of computational variables governing the adaptive belief updating. The observed changes in belief updating and arousal dynamics may account for the emergence of cognitive biases in psychotic psychopathology. Our results open a new window on the pathophysiology of mental disorders.

## Introduction

Many decisions entail the protracted accumulation of noisy information about the state of the world – a process referred to as belief updating. Dynamic belief updating is key for healthy cognition, particularly in environmental contexts characterized by multiple sources of uncertainty. Aberrant belief updating can produce biased representations of these environments and, consequently, false expectations about future events. Such problems are likely to result in maladaptive reasoning and decision-making as is evident in many psychiatric disorders including psychosis.

The possibility of (often hidden) change of environmental state is a key source of uncertainty in natural environments ^1^, and incorporating this feature into laboratory decision-making tasks has provided new insight into the mechanisms of belief updating ^2^. Human participants seem to approximate normative belief updating in a variety of change-point tasks ^2–7^. The normative process entails a dynamic ‘upweighting’ of new evidence at moments when the probability of a state change (‘change-point probability’, CPP) or uncertainty in the belief are high ^2,5,8,9^. Such transient increases in evidence sensitivity are, in part, mediated by phasic arousal responses, measured through the dilation of the pupil ^2,5,8^, an established marker of central arousal state_10-15._

The above insights may have important implications for understanding psychosis. Problems in constructing accurate beliefs are a hallmark of psychotic experiences, such as delusions (i.e., fixed beliefs contradicting the evidence) or hallucinations (non-veridical percepts). While these experiences are key symptoms of patients diagnosed with schizophrenia, they are also reported by people in the general population ^16^. An emerging line of research points to aberrations of probabilistic reasoning in psychosis ^17–21^. One aberration is the tendency to ‘jump to conclusions’ – a premature commitment to a particular hypothesis without assessing all the available information ^22,23^. A related phenomenon is the so-called ‘bias against disconfirmatory evidence’ – a failure to change an initial hypothesis about a hidden state in the face of evidence against that hypothesis ^24,25^. Such biases may result from alterations of evidence weighting, giving too strong a weight to initially encountered observations in determining belief states.

Another, and so far disjunct, line of research has established a dysregulation (typically, increase) of tonic arousal levels in psychosis ^26–30^. There is an opponent interplay between tonic and phasic activity modes of central arousal systems ^31^. Thus, increased tonic arousal levels observed in psychosis may reduce the phasic, pupil-linked arousal responses tracking CPP or uncertainty during belief updating. This, in turn, may reduce the dynamic evidence re-weighting in changing environments, altering the belief updating process overall.

Here, we developed an integrated approach to test these hypotheses. We relate individual differences in the proneness to psychosis to adaptive belief updating profiles in a volatile environment, as well as to the associated pupil-linked arousal responses. We first established that the mechanistic insights gained in previous work ^2,5,7^ generalize to a representative community sample. We went on to show highly specific aberrations in evidence weighting profiles and inference-driven pupil responses in individuals prone to psychosis. Finally, we established a close correspondence between individual evidence weighting profiles and inference-driven pupil responses.

## Results

We related individual differences in adaptive belief updating and pupil-linked arousal responses to the individual psychosis proneness in a large community sample of 90 participants (Figure 1; see Methods for details on recruitment), combining the behavioral task (Figure 1a), modelling (Figure 1b) and psychophysiological (Figure 1c) approach developed in Murphy et al. ^2^ with responses to an established self-report questionnaire.

**Figure 1.**
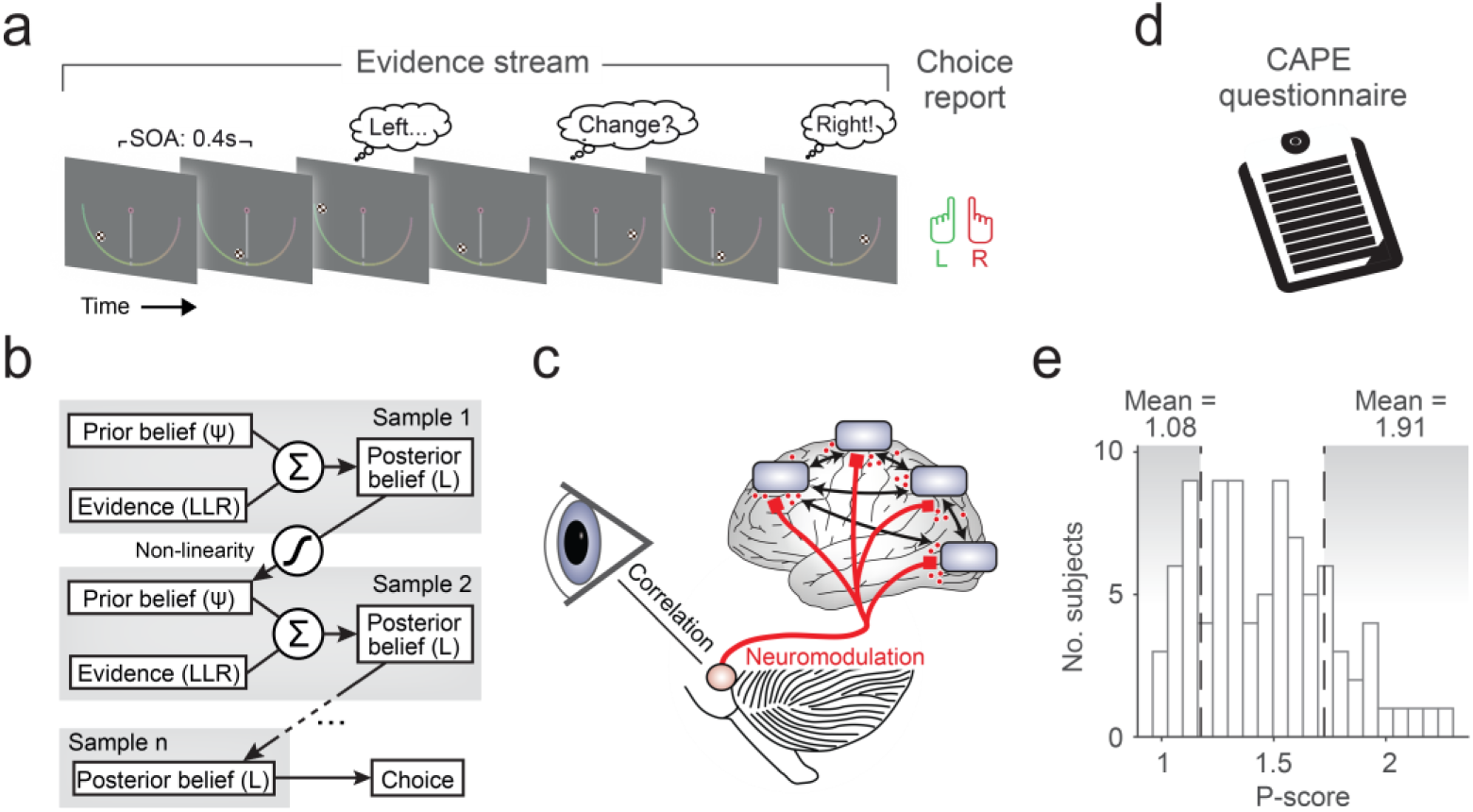
Behavioral task and approach. (**a**) Schematic of two-alternative perceptual choice task with hidden state changes. Locations of successive evidence samples (checkerboard patches) were drawn from one of two noisy sources that switched unpredictably over time, and participants reported the inferred active source when the sequence terminated. Each possible sample location provided a unique amount of evidence for one or the other alternative, as illustrated by the color bar shown to participants throughout (yellow ≈ no evidence, green ≈ strong evidence for leftward source, red ≈ strong evidence for rightward source). The sample stimulus-onset asynchrony (SOA) was 400 ms. (**b**) Schematic of normative belief updating process. (**c**) Tracking central arousal state through monitoring of pupil diameter. Rapid, non-luminance-mediated dilations of the pupil are an established proxy of the activity of neuromodulatory brainstem systems with wide projections throughout the brain, by which they control cortical network state. See main text for details. (**d**) Measuring individual psychosis proneness via CAPE questionnaire. (**e**) Distribution of psychosis proneness (P-scores) extracted from questionnaire data. Vertical dashed lines indicate cutoffs for lowest and highest P-score quintiles, determining subgroups used for the majority of our analyses; means are mean P-scores within each subgroup. **a**, **b** Adapted from ref. ^2^. **c** Adapted from ref. ^11^.

Psychosis proneness was assessed with the frequency dimension of the positive (P) score of the Community Assessment of Psychic Experiences CAPE; ^32,33;^ Figure 1d. This score was chosen because it assesses the type of psychotic experiences with an intuitive relationship to aberrant belief formation (i.e. hallucinatory experiences and delusional beliefs), is most indicative of psychosis proneness across the continuum and can be used as a screening tool to identify those at risk of future development of psychosis ^34,35^. In line with the recommended scoring procedure, the positive score (henceforth: P-score) was calculated by averaging responses across a subset of 20 predefined questionnaire items to compute a single summary metric. We found a broad range of P-scores across our sample, reaching values found in individuals fulfilling established criteria for ultra-high risk of developing psychosis P-score, frequency dimension > 1.7; ^35^ in a modest (20%) fraction of participants (Figure 1e; see Supplementary Figure 1 for distributions of the N (negative) and D (depressive) scales).

The task (Figure 1a) required accumulation of evidence provided by discrete visual ‘samples’, under the possibility of hidden changes in the underlying (categorical) environmental state that participants had to infer. On each trial, between 2 and 10 evidence samples were presented sequentially, with the spatial location of each sample providing noisy information about the hidden state. Sample locations were generated from one of two probability distributions (one per environmental state). The generative state was chosen randomly at the beginning of each trial and could then change after each sample, with a fixed probability (task hazard rate, *H*) of 0.1 in the main experiment. Participants were asked to report the generative state at the end of each sequence (left-or right-handed button press).

The normative computation maximizing accuracy on this task (Figure 1b) entails the accumulation of evidence samples (expressed as log-likelihood ratios, *LLR*, quantifying strength of support for one over the other state) into an evolving belief *L* (also expressed as log-odds ratio) that governs the final decision ^4^. The key difference between the normative model and standard evidence accumulation models (such as drift diffusion ^36^) lies in a non-linear function through which the updated belief after each sample *n* (*L_n_*) is passed to compute the prior for the next updating step (*ψ_n+1_*). This non-linearity depends on the agent’s estimate of the environmental volatility, the subjective hazard rate ^*H*^^. This way, the normative model strikes an optimal balance between formation of strong beliefs in stable environments versus fast change detection in volatile environments. In the following, we used this normative model in its pure form (called ‘ideal observer’ below, i.e., with exact knowledge of the task *H* and without any internal noise) as a benchmark against which to compare the behavior of human participants. We also fitted variants of this model combined with deviations from this pure form to each participant’s choices.

### Participants implement adaptive belief updating with non-linear evidence weighting

Complementary model-based behavioral analyses indicated that most participants approximated the normative belief updating strategy, subject to some internal noise and misestimation of the task hazard rate (Figure 2). While, as expected, all participants achieved accuracies below those of the benchmark provided by the ideal observer, most performed better than alternative, sub-optimal strategies (‘perfect integration’, that is uniform evidence weighting and no information loss; and a ‘choose based on last sample’ heuristic) even when these were not contaminated by any noise (Figure 2a). The participants’ choices were also more consistent with those of the ideal observer (same choice on 85.7 ± s.e.m. = 0.45% of trials) compared to the alternative strategies (perfect integrator: 78.1 ± 0.38%, *P*<0.0001; last-sample heuristic: 76.4 ± 0.62%, *P*<0.0001; paired-samples permutation tests).

**Figure 2.**
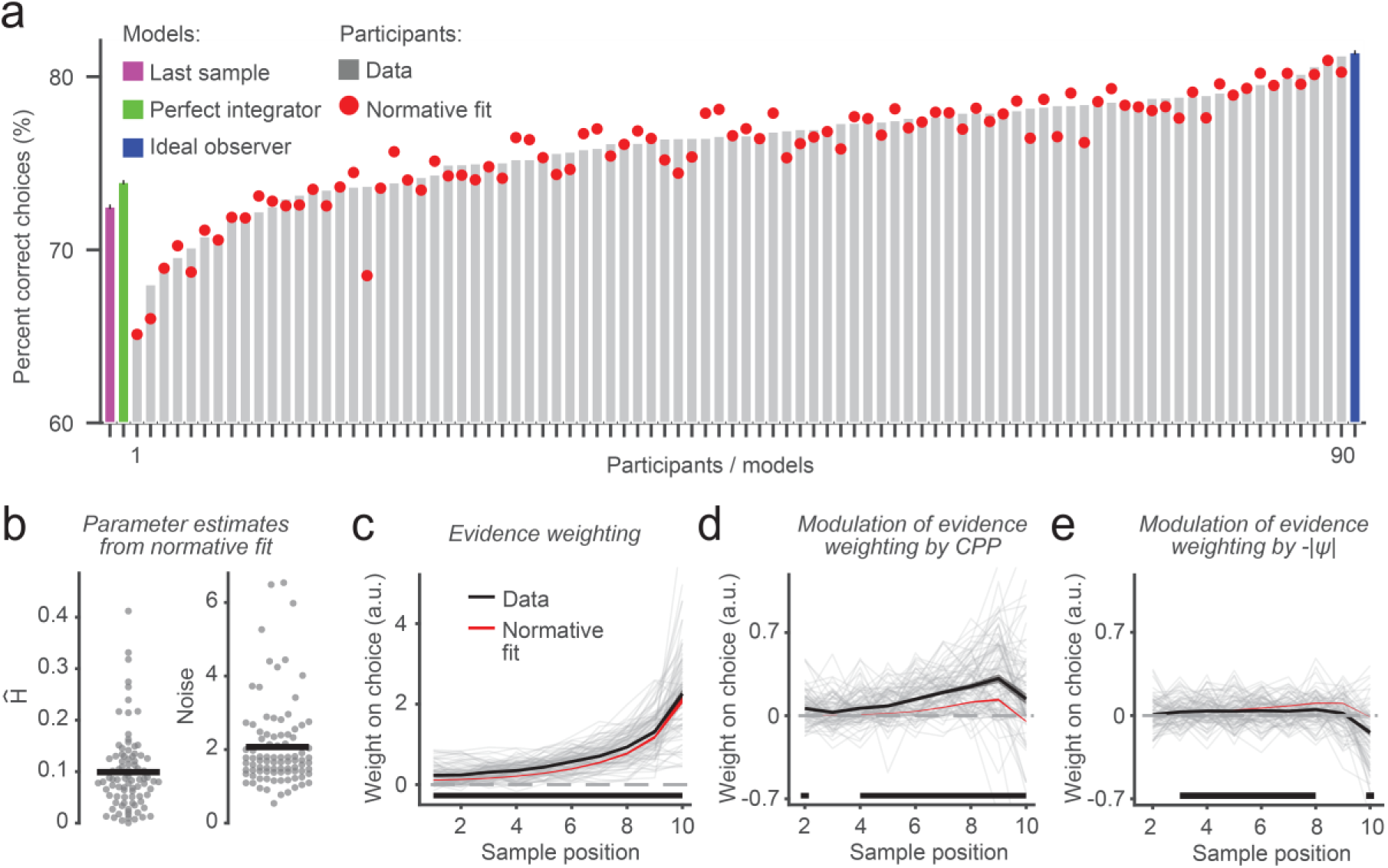
Behavioral signatures of adaptive belief updating. **(a)** Choice accuracies of n=90 human participants (gray bars), with ideal observer (navy), perfect integrator (green) and deciding based on last sample heuristic (green) shown for reference. Red dots, accuracies of normative model fits. (**b**) Best-fitting parameters of normative model fits for individual participants (grey circles) and group means (black horizontal bars). **(c, d, e)** Time-resolved weight of evidence on choice (**c**), and its modulation by change-point probability (CPP, **d**) and uncertainty (-|ψ|, **e**). Black, mean and s.e.m. (shaded areas) of participants’ data; significance bars, time points where weights differ from zero (p<0.05, two-tailed cluster-based permutation test; evidence weighting and largest CPP and -|ψ| modulation clusters, p<0.0001). Red shadings, fits of normative model. Gray thin lines, individual participants.

The notion that participants approximated normative belief updating was also supported by fitting alternative variants of the normative model to their choices (Figure 2; Supplementary Figure 2). Informed by model validation and comparison (Supplementary Figure 2; *Methods*), the variant of the normative model that we used for all further analyses contained two free parameters: a subjective hazard rate (^*H*^^) that could deviate from the task hazard rate, and a decision noise parameter Figure 2b; ^2,4,7^. The choices of this best-fitting model variant (‘normative fit’ in Figure 2a, red) were highly consistent with participants’ choices (86.9 ± 0.42%). The model fits also suggested that while at the group level participants appeared to form accurate estimates of the level of volatility in the task (^*H*^^ = 0.099 ± 0.008 compared to task *H* of 0.1; *P*=0.9, permutation test), there was significant variation in these estimates across individual participants (^*H*^^ range = [0.0003, 0.412]; Figure 2b). Overall, these results are in line with previous work in smaller, and less representative, samples ^2,4,7,37^.

Critically, our current participants also exhibited the same average evidence weighting profiles (Figure 2c-e) that were previously identified as diagnostic of the normative accumulation process and observed in smaller, less representative, samples – specifically the dynamic modulation of the impact of evidence on choice by change-point probability (*CPP)* and uncertainty -|ψ|; ^2^. *CPP* is the posterior probability of a change in generative state, given the subjective hazard rate, prior belief state and new evidence sample. Because it is contingent on internal belief states and representations of environmental volatility, this latent variable captures a high-level form of surprise. Uncertainty is captured by the (sign-flipped) absolute strength of the belief before a new sample is encountered (i.e. the prior). We estimated *CPP* and -|*ψ*|, for each sample *n*, from belief trajectories *L_n_* predicted by the normative fits Methods; ^2^, and quantified the evidence weighting profiles through logistic regression.

The first set of regression weights quantified the direct leverage of *LLR* on choice ’psychophysical kernel’; ^38,39^. These weights provided evidence for temporal integration (all weights significantly above 0; Figure 2c) with a clear recency effect, i.e., stronger impact on choice for late versus early samples (*P*<0.0001, two-tailed permutation test on ‘kernel difference’ of the weights for first 5 *vs*. last 5 samples).

The second and third sets of regression weights captured the modulation of evidence weighting by *CPP* and -|*ψ*|, respectively (Methods). The *CPP* modulation can be interpreted as the readiness to change one’s belief state in the face of contradictory (disconfirmatory) evidence, a construct prevalent in psychosis research (see Discussion); while the -|*ψ*| modulation captures a tendency to give greater weight to new information when in a state of uncertainty, often an adaptive policy in changing environments ^40,41^. Both modulations reflect the dynamic, context-dependent weighing of new information that permits the flexible belief updating prescribed by the normative model ^2^. There was a robust modulation of evidence weighting by *CPP* (*P*<0.0001 for cluster encompassing sample positions 4-10, cluster-based permutation test; Figure 2d), particularly strongly for samples in the second half of the evidence stream (*P*<0.0001, two-tailed permutation test on *CPP* ‘kernel difference’). There was also a reliable -|*ψ*| modulation (*P*<0.0001 for cluster encompassing sample positions 3-8; Figure 2e). As in previous investigations in similar task contexts ^2^, we found that the effect of *CPP* was the stronger of these adaptive modulations of evidence weighting (*P*<0.0001, two-tailed permutation test). As also reported previously ^2^, the observed modulation of evidence weighting by *CPP* was stronger in the participants than the normative model fit to their choices (Figure 2d, compare black and red).

In sum, participants based their decisions on an approximately normative belief updating process, subject to internal noise and biased encoding of the environmental hazard rate. This replicates previous findings ^2^, here in a community sample of participants more representative of the general population than the samples used in previous studies of the computational and neural bases of belief updating in changing environments but see ^6^.

### Participants prone to psychosis exhibit altered signatures of belief updating

While the previously discovered evidence weighting effects were statistically significant at the group level, we also observed substantial individual differences in the evidence weighting profiles (Figure 2c-e). We related these individual differences to those in psychosis proneness as indicated by the P-score of each participant (Figure 1e). In order to evaluate the detailed shapes of the weighting profiles, we grouped the individual kernels into bins (quintiles) defined by the associated P-scores and focused our comparisons on the two participant subgroups with the lowest and highest scores (Figure 3; see also Supplementary Figure 3). The mean scores were 1.08 (s.d. = 0.057) for the ‘low P subgroup’ and 1.91 (s.d. = 0.163) for the ‘high P subgroup’. The latter corresponds well to the mean P-score obtained in individuals fulfilling the criteria for ultra-high-risk status for psychosis 1.9, CI = 1.71–2.02; ^35^. We also computed continuous correlations between scalar indicators derived from the kernels (as well as other measures of behavior) and the individual P-scores (Supplemental Figures 4 and 5).

**Figure 3.**
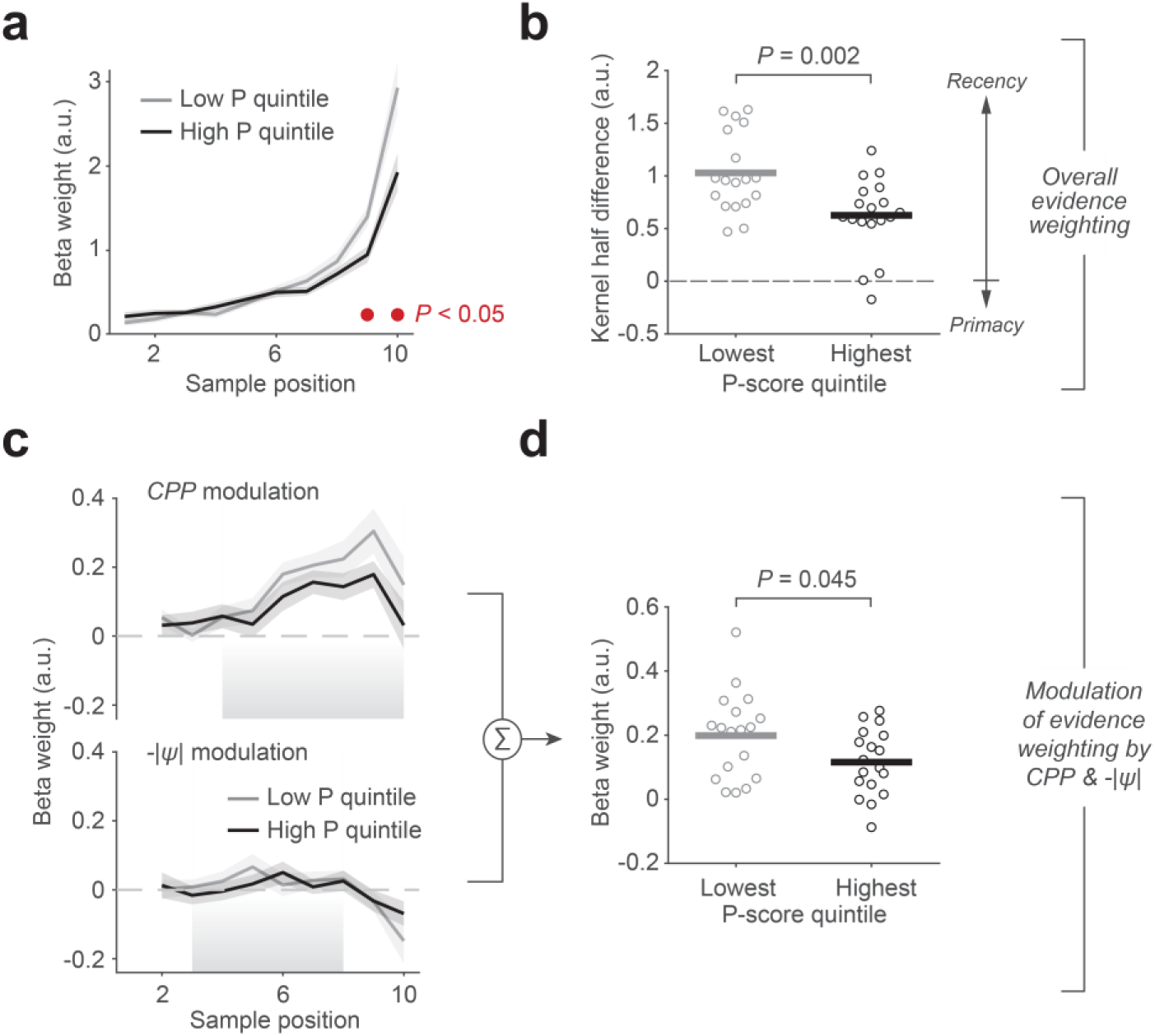
Change in belief updating profiles for individuals with high psychosis proneness. **(a)** As Figure 2c, but now for subgroups of participants with the lowest (grey) and highest (black) quintiles of P-scores. (**b**) Kernel half-difference summary measure (subtraction of mean weighting of first 5 samples from mean weighting of last 5 samples) capturing degree of recency in evidence weighting, plotted for both lowest and highest P-score quintiles. (**c**) As Figure 2d,e, but now for subgroups with lowest and highest P-score quintiles. (**d**) Summary measure capturing summed strength of modulations of evidence weighting by CPP (mean modulation weights over sample positions 4-10, significant cluster in Figure 2d) and -|ψ| (mean modulation weights over sample positions 3-8, significant cluster in Figure 2e), plotted for lowest and highest P-score quintiles. Horizontal lines in **b** and **d**, mean of data from each subgroup; circles, individual participants. P-values, two-sample permutation tests (two-tailed).

There was a clear relation between the P-scores and the model-estimated noise parameter (Supplemental Figure 4d,h) and mixed evidence for an association with subjective hazard rate^*H*^^ (Supplemental Figure 4c,f). Further, participants’ overall behavioral performance – evaluated as accuracy, and consistency of choices with those of the ideal observer, specifically on trials containing at least one change point – was better in the low P subgroup than the high P subgroup (Supplemental Figure 4a,b), albeit without a robust effect for the corresponding continuous correlations (Supplemental Figure 4e,f). These observations, along with the inferred high-risk status of the well-isolated high P subgroup, motivated our main focus on the comparison between the two subgroups in our analysis of evidence weighting profiles as well as pupil dynamics.

The high P subgroup gave overall less weight to evidence arriving late in the evidence stream, as quantified in terms of the individual sample comparisons (Figure 3a) as well as the strength of the above-described overall recency effect (i.e., kernel half difference, Figure 3b). Furthermore, this subgroup exhibited a generally weaker adaptive component to their belief updating, reflected in smaller summed weights of the *CPP* and -|*ψ*| modulations identified above (Figure 3c,d). This effect was significant when summing regression coefficients across both variables (Figure 3d) and marginally significant for *CPP* alone (Supplementary Figure 5). Similar effects were observed for the kernel half difference, but not for the summed or individual modulation effects, in the continuous correlations across all subjects (Supplementary Figure 5).

In sum, we found differences in the adaptive, non-linear belief updating process between individuals scoring on the lowest end of the psychosis continuum, and those with scores approximating those of individuals who fulfill established criteria for being at ultra-high risk of developing psychosis ^35^. These alterations were in line with a stronger ‘stickiness’ of belief states formed early on during processing of evidence streams in the high P subgroup.

### Participants prone to psychosis also have altered pupil dynamics

Previous work has shown that phasic arousal encodes *CPP*, -|*ψ*| and related variables and, at least in part, mediates their impact on learning rate and evidence weighting ^2,5,8,42^. We isolated phasic arousal responses to individual samples in terms of the first temporal derivative of the pupil diameter signal measured during task performance ^2,43^ (Figure 4a). Pupil diameter is as an established proxy for changes in arousal state (McGinley et al, 2015; Joshi & Gold, 2020). We focused on its first derivative in order to increase temporal precision and specificity for noradrenaline release ^15^. We then regressed this rapid response component on single-sample *CPP* and *-IψI* estimates in a time-resolved fashion (Methods), without making assumptions about the generator mechanism ^44^. This approach revealed robust pupil encoding of both computational variables (Figure 4d), again replicating previous findings ^2^.

The evoked pupil responses also differed between the high and low P subgroups (Figure 4b-d). The overall responses during the trial were reduced in high P individuals, when quantified as either (i) evoked responses in the raw pupil signal and averaged across the trial epoch (Figure 4b; marginally significant) or (ii) as the response of the pupil derivative (Figure 4c) evaluated in an early time window that contained the largest excursion of the derivative response (Figure 4a, dashed line). Furthermore, the encoding of belief updating variables in the pupil derivative was less reliable in the high P subgroup (Figure 4e; Supplementary Figure 6).

**Figure 4.**
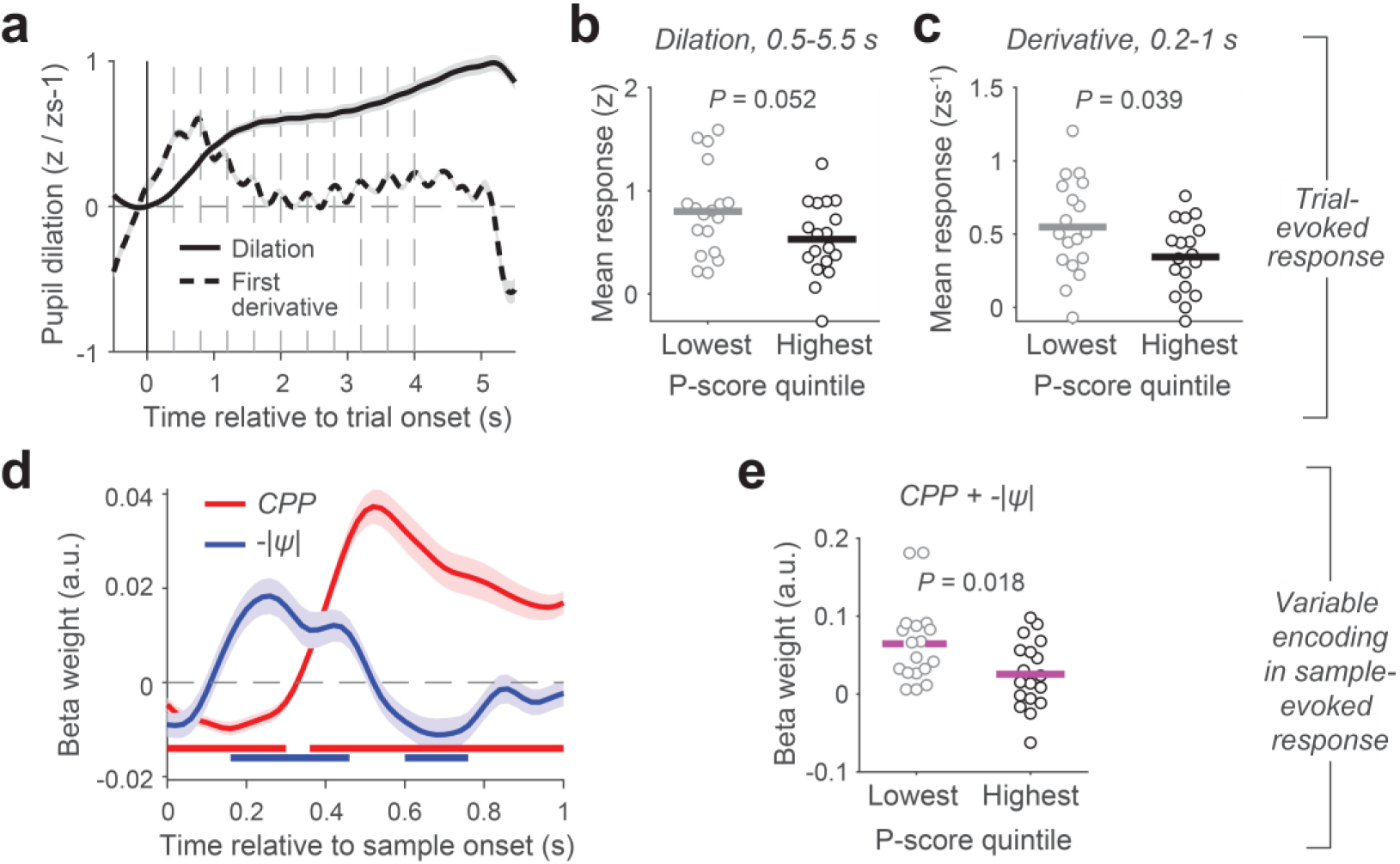
Evoked pupil responses and individual differences. (**a**) Average trial-related pupil response (solid line) and its first derivative (dashed line). Dashed grey vertical lines, sample onsets. (**b, c**) Both measures of overall trial-evoked response for the low and high P-score subgroups. (**d**) Encoding of change-point probability and uncertainty in pupil responses evoked by individual evidence samples. (**e**). Encoding of change-point probability and uncertainty in pupil responses (pooled) for different P-score subgroups. Shaded areas in panels **a**,**d** indicate s.e.m. Significance bars in **d**, p<0.05 (two-tailed cluster-based permutation test). Horizontal lines in **b,c** and **e**, mean of data from each subgroup; circles, individual participants. P-values, two-sample permutation tests (two-tailed).

In sum, not only did the high P subgroup exhibit altered belief updating dynamics, but their task-related recruitment of pupil-linked arousal was weaker, both in terms of the overall evoked response and sensitivity to the computational variables that modulate the belief updating computation.

### Inference-related pupil dilations predict individual evidence weighting profiles

Given that high and low P subgroups differed in both their above-described behavioral signatures of adaptive evidence weighting (i.e., kernel metrics, Fig. 3b,d) and the encoding of computational variables from the inference process in their pupil responses (specifically, *CPP* and uncertainty Fig. 4e), an obvious question is how the individual differences in these variables related to one another. Indeed, we observed robust correlations between the pupil sensitivity to *CPP* and uncertainty, and both evidence weighting signatures: the kernel difference (Figure 5a) and the (pooled) modulation of evidence weighting by *CPP* and uncertainty (Figure 5b). Weaker relationships to the behavioral evidence weighting measures were present for the overall task-evoked pupil response (magnitude or derivative) across the entire trial (Supplementary Figure 7a, b). Taken together, these results show that the individual level of recruitment of pupil-linked arousal by inference-related variables reflected individual differences in adaptive evidence weighting.

**Figure 5.**
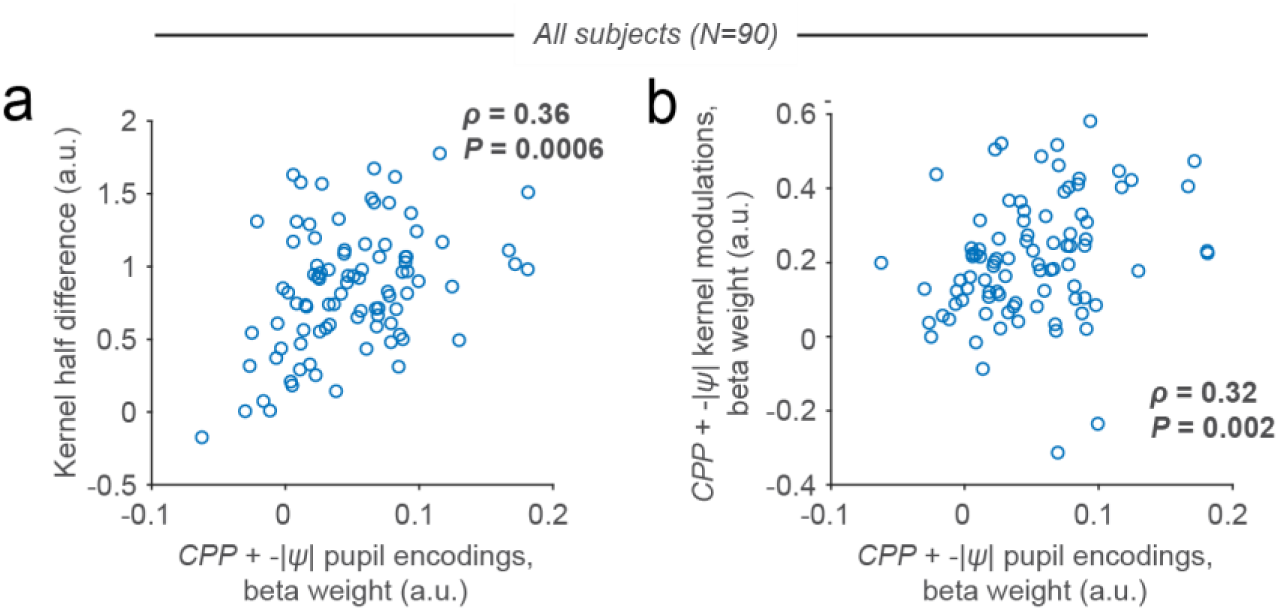
Relationship between individual belief updating and pupil metrics. Scatterplots show across-subjects correlation between encoding of CPP and -|ψ| in sample-related pupil responses (pooled) and the two evidence weighting (kernel) metrics from Fig. 3b,d: specifically, variable encoding in pupil response plotted against (**a**) kernel half-difference and (**b**) summed CPP and -|ψ| modulation weights. Circles, participants. Correlation coefficients and P-values, Spearman correlation. See Supplementary Figure 7c,d for the correlations evaluated separately for CPP and -|ψ|.

### No effect for other clinical scores measuring negative and depressive symptoms

Our analyses focused on the P-score of the CAPE questionnaire because of the rationale underlying the study. Indeed, the relationships with belief updating profiles and pupil dynamics reported above were specific to the P-score, with no effects for the other two scores measuring negative (N) and depressive (D) symptoms, respectively (Supplementary Figure 8).

### No relationship between psychosis proneness and working memory performance

Many studies into the pathophysiology of schizophrenia have focused on working memory ^45,46^, the ability to hold information online for several seconds and manipulate it for the control of behavior ^47,48^. Deterioration of working memory is consistently associated with schizophrenia ^49^, in particular with the negative subdomain ^50^, and may provide early cues for the development of the disorder ^51^. For these reasons, and to test for the specificity of the effects observed in our belief updating task, we also related the CAPE questionnaire scores to participants’ performance in a classical delayed match-to-sample working memory task (Methods). We observed no reliable associations between working memory task performance and scores on any of the three scales that comprise the CAPE (Supplementary Figure 9). This indicates that the relationship between psychosis proneness and adaptive decision-making established above is driven by individual differences in the dynamic inference process, rather than by a possible contribution of working memory integrity to performance of our belief updating task ^52^.

## Discussion

Our work sheds new light on the mechanisms underlying individual differences in belief updating and their relationship to latent psychopathology. To this end, we built on recent computational and physiological insights into group average behavior on the same task as the current one ^2^. This previous work established that (i) participants approximated the normative belief updating strategy for this task, (ii) the underlying belief state was selectively encoded in the slow dynamics of action plans in parietal and frontal cortical regions, (iii) the computation of this belief state emerged from recurrent interactions in local microcircuits equipped with attractor dynamics, and (iv) pupil-linked phasic arousal responses contributed to the dynamic upweighting of evidence samples associated with high change-point probability and uncertainty, a process that was linked to modulation of the evidence encoding in visual cortex. Critically, this prior work also (v) identified diagnostic signatures of the non-linear, adaptive belief updating process (and underlying circuit dynamics) in the form of evidence weighting profiles ^2^.

Here, we show that the normative belief updating process also accounted for the behavior of a large community sample overall, and that this sample exhibited previously established diagnostic signatures of adaptive belief updating as well as encoding of key computational variables entailed in this process in their evoked pupil responses. Critically, we discovered that a subset of individuals prone to psychosis tended to give less weight to evidence that arrived late in the evidence stream and exhibited weaker combined dynamic modulations by change-point probability and uncertainty. These signatures point to an overall stronger persistence of initially formed belief states, in line with formation of cortical attractor states that are more resistant to subsequent change ^2^. Furthermore, this tendency was associated with smaller task-evoked pupil responses and less precise encoding of change-point probability and uncertainty in the pupil-linked arousal responses to individual samples.

Delusions and hallucinations are prominent symptoms of schizophrenia but are also reported by people in the general population in an attenuated form. This has led to the notion of a psychosis continuum ^16^ ranging from people who report never to have experienced even the mildest type of psychotic experience to those diagnosed with schizophrenia. Located in between these two extremes are various degrees of risk states characterized by subclinical psychotic experiences of varying frequency and intensity. Self-report measures of psychotic experiences, such as the CAPE questionnaire employed here ^32^, are commonly used to estimate a person’s position on this continuum, with higher scores indicating higher proneness to psychosis. The CAPE has been extensively validated to this end ^32,34,35,53,54^ and shows good test-retest reliability ^32^. The self-reported dimensions of psychosis that it purports to assess are associated with the corresponding interview-based dimensions ^32,55^. The P-scores that we focused on are suited to screen for psychosis risk ^34,35^ and for clinical psychosis ^53^. The 3-dimensional structure is stable across different populations and language versions, including the German Version used here ^54,56^. We, therefore, conclude that the self-reported dimensions of psychotic experiences assessed here are reliable and valid.

Our findings highlight the importance of the detailed, model-guided quantification of evidence weighting profiles (so-called psychophysical kernels) that we have derived computationally and validated physiologically ^2^. These profiles can identify even nuanced and idiosyncratic alterations in the dynamic weighting profiles, which may not be fully captured by parameter estimates from model fits. We found a consistent effect of individual P-scores on decision noise and a weak effect on the subjective hazard rate (depended on inclusion or exclusion of an outlier). In line with recent insights from a learning task with change-points applied to schizophrenia ^57^, this pattern of results indicates that static parameter estimates can fail to capture important deviations in belief updating dynamics. Such deviations are instead well captured by the psychophysical kernels that we here estimated for each individual.

Our results are broadly consistent with predictive coding accounts of psychosis ^58–61^. Specifically, the reduced pupil encoding in the high-P subgroup of the inconsistency between prior and evidence (as measured by change-point probability) that we observed here suggests an impaired computation of cognitive prediction errors, which other work has linked to schizophrenia and delusion severity ^62^. Furthermore, it is tempting to relate our observation of stronger stickiness of initially formed beliefs to the notion of overly strong predictions about a higher-level context as a source of false beliefs ^63^. Our results also support accounts of aberrant probabilistic reasoning in psychosis that focus on the over-expression of two cognitive biases: jumping to conclusions ^22,25^ and bias against disconfirmatory evidence ^24,25^. Both biases are assumed to contribute to the formation of false beliefs that are the essence of delusions ^64–68^.

Importantly, the above accounts of biased probabilistic reasoning are mainly based on behavioral evidence from versions of the so-called beads task without hidden state changes ^69,70^, and on analyses that suffer from interpretational limitations. First, the neural bases of the cognitive computations underlying performance of these tasks is unknown. Second, inferences about biases in this and similar tasks are indirect, through latent model variables or substitute measures of behavioral performance; the dynamics of belief updating are not probed as directly through analysis of evidence weighting profiles (psychophysical kernels) widely used in decision neuroscience ^2,71^. Third, in the classical ‘draws to decision’ version of the beads task ^69,70^, the subject draws beads from one of two jars with fixed (and already learned) probabilities until committing to a decision about which of the two jars they are drawing from. This version requires the setting of decision bounds to determine how much evidence is accumulated before the subject decides to stop accumulating and make a response. This may conflate biases in evidence accumulation *per se* with a general proneness to commit to decisions and/or to stop engaging in the task. Fourth, and most fundamentally, the standard tasks that have been used in this literature have lacked hidden state changes, which are an essential feature of natural environments ^1^. Indeed, our recent analyses indicate that the presence of such hidden state changes in laboratory tasks is critical for understanding the significance of non-linearities of human belief updating and the underlying cortical (attractor) dynamics ^2^.

Our current model-based behavioral and pupillometry approach overcomes these limitations. We used an evidence accumulation task with hidden change-points, which provided direct access to the dynamics of evidence weighting and the neurophysiological basis which we have previously characterized in detail. This enabled us to precisely quantify two biases: (i) Primacy vs. recency in temporal evidence weighting and (ii) modulation of evidence weighting by the contextual variables change-point probability (which depends on the inconsistency of a new evidence sample with the current belief state) and, to a weaker extent, uncertainty. Because the number of samples presented on each trial are under the experimenter’s (rather than the subject’s) control, these biases are independent of stopping rules and directly inform about belief updating. Stronger tendency toward primacy in evidence weighting can be interpreted as a stronger tendency toward jumping to conclusions ^72^. Likewise, a reduced modulation of evidence weighting by CPP indicates a bias against disconfirmatory evidence. Both effects were present (the latter trending when isolated from the uncertainty modulation) in subjects with the highest proneness to psychosis. Finally, and critically, our approach enabled linking the behavioral identification of these biases to specific, inference-driven components of task-evoked pupil responses.

Our results open a window on the pathomechanisms of psychosis. We observed that precision of the encoding of CPP and uncertainty in pupil responses, a high-level, cognitive component of the pupil dynamics in our task, was particularly closely related to individual evidence weighting signatures and was also related to delusion proneness. It is tempting to speculate that the previously established hyper-arousal in psychosis renders the arousal system less responsive to high-level variables computed in the process of belief updating, specifically CPP and uncertainty. This reduces the dynamic upweighting of internal representations of the momentary evidence observed in sensory cortex ^2^, translating into a smaller than required weight of that evidence in the updating of the belief state in downstream cortical regions. As a consequence, an individual’s beliefs will become ‘sticky’. Our findings were obtained in an emotionally neutral belief updating task operating on short timescales and unrelated to the common (often social) contents of psychotic beliefs. It is all the more striking that specific computational measures of behavior and arousal response derived from this kind of task context relate to overall psychosis proneness. This suggests that the pathophysiology of psychosis may affect not only high-level (social, emotional) reasoning that dominates delusions, but also inferences about low-level (and emotionally neutral) properties of the sensory environment.

We conclude that aberrations in belief updating and arousal dynamics are evident in individuals who have moderate-to-high levels of self-reported psychosis proneness but have not been clinically diagnosed with schizophrenia. These aberrations may give rise to psychotic symptoms (i.e., delusions), by increasing the likelihood of holding on to erroneous beliefs. They may also serve as a risk marker of psychotic psychopathology. Our computational approach to understanding belief updating behavior and arousal may become a useful tool for future studies into the pathophysiology of psychosis and other mental disorders.

## Methods

### Recruitment and Sample

The study was approved by the ethics committee of the Faculty of Psychology and Movement Science at Universität Hamburg and included a total of 96 human participants. Given the skewness of P-scores in the population, with the majority scoring at the low end and few scoring at the higher end of the psychosis continuum, we oversampled participants in the high range aiming for 50% with a P-score above the 50th percentile within a large community sample ^54^. All participants provided written informed consent, were aged between 18-55 years old, fluent in the German language, had an IQ > 85 as assessed by the Multiple Choice Vocabulary Test ^73^, had normal/corrected-to-normal vision, were not pregnant, and had no dementia or other organic brain disorders, acute intoxication or acute suicidal tendencies. Participants received remuneration in the form of an hourly rate (€10 per hour), a bonus for completing both of two planned sessions (€15), and compensation if they incurred costs for a SARS-CoV-2-antigen test to participate in the study.

Four participants were excluded from all analyses due to failing to complete both testing sessions, and 2 more participants were excluded due to incomplete CAPE questionnaire responses. The remaining 90 participants (mean ± s.d. age of 31.6 ± 9.5 years, range 18-55; 48 females) were included in all analyses of data from the main behavioral task, having completed two experimental sessions (session 1: 180 min; session 2: 160 min). A further 2 participants were excluded from analyses of data from the delayed match-to-sample task having failed to complete any blocks of this task due to time constraints.

Of the 90 participants included in all main analyses, four self-reported a previous diagnosis of depression, one of emotionally unstable personality disorder (impulsive type), one of post-traumatic stress disorder (PTSD), one of obsessive-compulsive disorder (OCD), one of comorbid depression and attention-deficit hyperactivity disorder (ADHD), and one of psychosis due to substance abuse. The latter participant had a CAPE P-score of 2.1, which placed them in the highest P-score quintile. Except for the summed modulations of evidence weighting by *CPP* and -|*ψ*| (Figure 3d, trending at *P*=0.1 with this participant excluded), all effects reported in Figures 3 and 4 remained statistically significant even when this participant was excluded from the analysis.

For participants that completed the full experimental protocol, the first session consisted of study information and consent, brief intelligence screening via Multiple Choice Vocabulary Test ^73^ and visual attention and task switching task Trail-Making-Test (TMT) ^74^, a questionnaire battery, training on the main behavioral task and 7-8 experimental blocks of this task (see below). The questionnaire battery was administered after the first three experimental blocks, after which the remaining experimental blocks took place. The second session consisted of 8-9 blocks of the main behavioral task as well as training and between 2 and 3 experimental blocks of the delayed match-to-sample working memory task halfway through the set of blocks of the main task (see below). We also measured at-rest heart rate variability at the beginning of each session via electrocardiogram, though results pertaining to these data are not reported here.

### Questionnaire battery

The Community Assessment of Psychic Experiences (CAPE) ^32,33^ is a self-report questionnaire consisting of 42 items that assesses frequency and distress of lifetime psychotic experiences. It subdivides into three factors: negative symptoms, positive symptoms, and depression symptoms. Previous research has shown that the CAPE has strong evidence of both convergent and discriminative validity, as demonstrated by Hanssen et al. ^55^. Additionally, the instrument has been shown to be reliable over time, with good test-retest reliability ^32^. The German version of the CAPE, which we employed here, has also been validated in terms of its factorial and criterion validity ^54^. Here, we focused on the frequency dimension of the P-score, both in regard to sampling the high end of the continuum and in terms of the main analyses. The D-and N-scores were used to assess specificity of the findings.

The Trier Inventory for Chronic Stress (TICS) ^75^ and Trauma History Questionnaire (THQ) ^76^ were used to assess the chronic stress load of the participants. Self-efficacy was assessed with the General Self-Efficacy Scale (SWE) ^77^ and achievement motivation with the Achievement Motives Scale-Revised (AMS-R) ^78^. Due to the fact that these scales were utilized to address hypotheses unrelated to the present study, we do not describe them in detail here nor report related results.

### Main behavioral task

The main task was a two-alternative forced choice task in which the generative task state *S*={*left*, *right*} could change unpredictably. Participants were asked to maintain fixation at a centrally presented mark throughout the trial, monitor a sequence of evidence samples and report their inference about *S* at the end of the sequence.

Stimuli were generated using Psychtoolbox 3 for Matlab ^79^. Visual stimuli were presented in a behavioral laboratory on a Dell P2210 22-inch monitor with resolution set to 1680 x 1050 and refresh rate to 60 Hz. Subjects were seated with their head in a chinrest 52 cm from the monitor during task performance.

Stimuli were presented against a grey background. Three placeholders were present throughout each trial: a light-grey vertical line extending downward from fixation to 7.4° eccentricity; a colored half-ring in the lower visual hemifield (polar angle: from -90 to +90° relative to bottom of vertical meridian; eccentricity: 8.8°) which depicted the *LLR* associated with each possible sample location; and a fixation mark as a black disc of 0.18° diameter superimposed onto a disk of 0.36° diameter with varying color informing participants about trial intervals. The colors comprising this half-ring and the fixation point were selected from the Teufel colors ^80^. Evidence samples consisted of achromatic, flickering checkerboards (temporal frequency: 10 Hz; spatial frequency: 2°) within a circular aperture (diameter = 0.8°), and varied in polar angle (constant eccentricity of 8.1°).

Samples were presented for 250 ms (sample-onset asynchrony (SOA): 400 ms). Samples were centered on polar angles *x_1_*,…,*x_n_* drawn from one of two truncated Gaussian distributions *p*(*x*|*S*) with variance *σ_left_*=*σ_right_*=27° and means symmetric about the vertical meridian (*μ_left_*=-17°, *μ_right_*=+17°). If a draw *x_i_* was <-90° (>+90°), it was replaced with -90° (+90°). *S* was chosen at random at the start of each trial and could change with a hazard rate *H*=*p*(*S_n_*=*right*|*S_n-1_*=*left*)=*p*(*S_n_*=*left*|*S_n-1_*=*right*)=0.1 after each sample. 65% of sequences contained 10 samples. The remaining 35% (randomly distributed in each block) contained (with equal probability) 2-9 samples and were introduced to encourage participants to attend to all samples. Each trial began with a variable fixation baseline period (uniform between 0.5-1.5 s) during which a stationary checkerboard patch was presented at 0° polar angle. This checkerboard then began to flicker, followed after 400 ms by the first evidence sample. The final sample of the sequence was then replaced by the stationary 0° patch and a ‘Go’ cue instructed participants to report their choice 0.7-1.2 s (uniform) after sequence end. They pressed left or right ‘CTRL’ keys on a keyboard, with their left or right index fingers, to indicate *left* or *right*, respectively. Auditory feedback 0.05 s post-response informed participants whether their choice corresponded to the true *S* at sequence end (see *Stimuli*): an ascending 350→950Hz tone (0.25 s) for correct choice and descending 950→350Hz tone for incorrect choice. This was followed by an inter-trial interval of 1.2 s when participants could blink. During the preparatory interval, sample sequence and subsequent delay, the color of the second fixation disk was light red; the ‘Go’ cue was this disk becoming light green; and, the inter-trial period was indicated by the disk becoming light blue.

Participants practiced the task prior to the main experiment, with detailed initial illustrations of the generative distributions and incremental increases in trial complexity. These instructions were delivered using a cover story that the task involved viewing sequences of footballs (the evidence samples) that two players kicked from the same point (fixation) but aiming at different goals (polar angles corresponding to the means of the two generative distributions). The players had quite bad aim (variance of the generative distributions) and could swap in and out of play between each kick with a fixed probability (the task *H*); and the participants’ task was to infer which player kicked the last football in each sequence. They were never informed about the exact task *H*, rather that the likelihood of a change in football player (*S*) from one sample to the next was “not very high. For example, there will be some trials in which there are no changes at all. However, many trials will have one change, and some trials can even have several changes.” Following training, participants completed 15-16 blocks of 86 trials of the task, spread over both testing sessions. They received feedback about their average performance at the end of each block, and were instructed to fixate the central disk and minimize blinking during the trial.

### Working memory task

Participants also completed a visuospatial delayed match-to-sample working memory task ^81^ presented on the same monitor as the inference task, again via Psychtoolbox 3. The task was to decide whether a sample stimulus and a test stimulus separated by a variable delay occurred in the same or different locations. Each trial began with presentation of a central white fixation cross (arm length: 0.8 degrees of visual angle, d.v.a.; arm thickness: 0.2 d.v.a.) that was present for the entire trial. After a variable baseline interval (uniform distribution with range 0.5-2.0 s), the sample stimulus was presented for 0.5 s, followed by the delay (1, 3 or 9 s, equiprobable) and then the test stimulus (0.5 s). Sample and test stimuli were circular checkerboard patches (diameter: 2.8 d.v.a.; spatial frequency: 1 cycle per d.v.a), appearing in the lower visual hemifield at a fixed eccentricity of 6 d.v.a.,. The sample could be presented at any of 12 equiprobable locations, ranging from ∼-76.15° to ∼76.15° of polar angle (fixed spacing ≈ 13.85°), while the most extreme samples could still be flanked by a ‘near non-match’ test stimulus on both sides (amounting to 14 possible test stimulus locations, spaced evenly from -90° to 90°). The test occurred at either the same location as the sample or at a different location (see below). Upon offset of the test stimulus, the fixation cross changed color from white to light blue, which prompted subjects to report their decision via right-or left-handed button press for “same” or “different” judgments, respectively. This response was soon (0.1 s) followed by visual feedback about its accuracy (“Correct” in green font; “Error” in red font; font size 36, presented 1.0 d.v.a. above fixation for 0.75 s). Each trial was followed by a fixed 2 s interval during which participants were instructed to blink if needed, and this was followed by the baseline period of the following trial.

The task was designed to consist of three trial categories, each with a desired frequency of occurrence within a block of trials: ‘Match trials’ (sample and test at identical positions, 1/3 of trials), ‘Near non-match trials’ (smallest possible sample-test distance of 13.85°; 1/3 of trials), and ‘Far non-match trials’ (sample-test distance randomly chosen from the remaining possible sample-test distances, which could be between 27.7° and ∼166.15° depending on the sample location; 1/3 of trials). Trials were presented in blocks of 63 trials each, within which the different delay durations and sample-test distances were randomly interleaved under the above-mentioned constraint. Subjects received feedback about their average performance at the end of each block. They were instructed to fixate the central cross and minimize blinking during the trial.

Subjects also underwent initial training to familiarize them with the task. This consisted of a general instruction of the task rules through PowerPoint slides, as well as practice with various aspects of the task (stimulus appearance, timings, response contingencies) that, as above, grew in complexity culminating in practice of full task trials. Once training was complete, each subject then performed 2 or 3 blocks of the task (mean = 2.25, s.d. = 0.44), with the specific number dependent on time constraints.

Trials on which participants pressed a key other than the designated response keys, response time was ≤ 0.2 s, or response time exceeded the participant’s mean response time by 4 s.d. were all excluded from analysis of response accuracy on the working memory task. Mean response accuracy was computed as the proportion of correct responses (i.e., “same” response on match trials and “different” response on non-match trials).

### Modeling of behavior

#### Normative model for main behavioral task

The normative model for the main task prescribes the following computation ^4^:

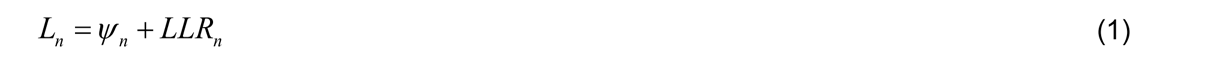

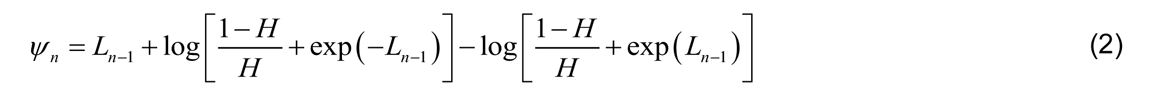

Here, *L_n_* was the observer belief after encountering the evidence sample *x_n_*, expressed in log-posterior odds of the alternative task states; *LLR_n_* was the log-likelihood ratio reflecting the relative evidence for each alternative carried by *x_n_* (*LLR_n_*=log(*p*(*x_n_*|*right*)/*p*(*x_n_*|*left*))); and *ψ_n_* was the prior expectation of the observer before encountering *x_n_*. We used this model to derive two computational quantities: *CPP* and -*|ψ|*. *CPP* was the posterior probability that a change in generative task state has just occurred, given the expected *H*, the evidence carried by *x_n_*, and the observer’s belief before encountering that sample *L_n-1_*, computed as follows (see ^2^ for derivation):

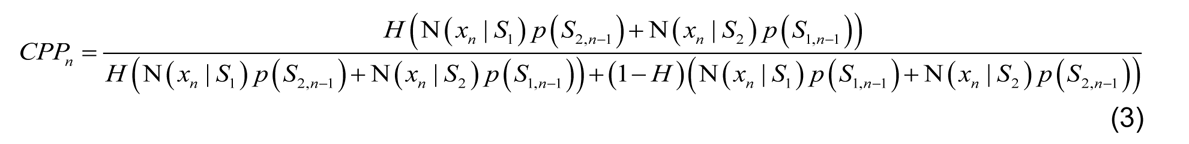

where N(*x*|*S*) denoted the probability of sample *x* given a normal distribution with mean *μ_S_* and s.d. *σ_S_*. Uncertainty was defined as the negative absolute of the prior (-*|ψ_n_|*), reflecting uncertainty about the generative state before observing *x_n_*.

#### Main task model fitting and comparison

We first computed the accuracy of participants’ choices with respect to the true final generative state, and compared this to the accuracy yielded by three idealized decision processes presented with identical stimulus sequences: normative accumulation (eqs. 1 and 2), perfect accumulation, and only using the final evidence sample. For each trial, choice *r* (*left*=-1, *right*=+1) was determined by the sign of the log-posterior odds after observing all samples: 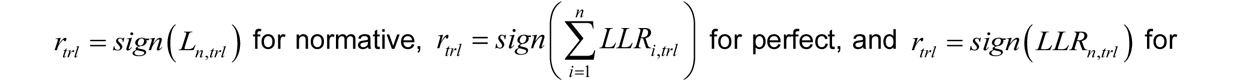 last-sample, where *n* indicated the number of samples presented on trial *trl*.

We fit variants of the normative model to participants’ behavior, assuming that choices were based on the log-posterior odds *L_n,trl_* for the observed stimulus sequence on each trial. In line with previous work with this model ^2,4^, different model variants had different combinations of the following free parameters: *L_n,trl_* was corrupted by a decision noise term *ν*, such that choice probability *r*^ was computed as:

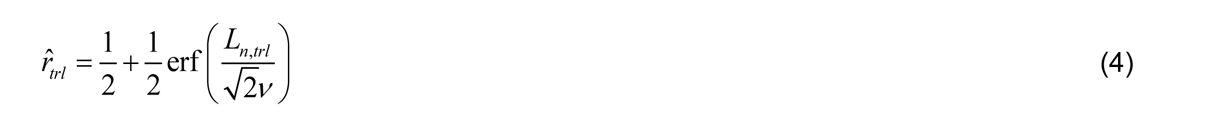

We also allowed for i) misestimation of the task *H* by fitting a subjective hazard rate parameter ^*H*^^; ii) a bias in the mapping of stimulus location to *LLR* such that subjective evidence strength *LLRn* = *LLR_n_* · *B* , where *B* captured the equivalent of under-(B<1) or over-estimation (B>1) of the signal-to-noise ratio of the task generative distributions (also known as ‘expected uncertainty’, ^9^); and iii) a bias in the weighting of evidence samples that were (in)consistent with the existing belief, such that *LLRn* = *LLR_n_* · *g* for any sample *n* where *sign (LLR_n_*) ≠ *sign (ψ_n_*).

We fitted the parameters by minimizing the cross-entropy between participant and model choices:

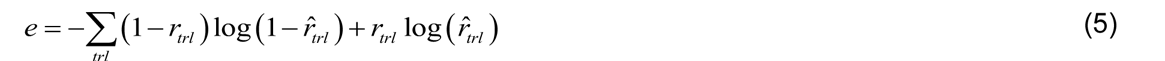

where *r_trl_* was the participant choice. This objective was minimized via particle swarm optimization (PSO toolbox) ^82^, setting wide bounds on all parameters and running 300 pseudorandomly-initialized particles for 1500 search iterations. The relative goodness of fit of different model variants was assessed by the Bayes Information Criterion (BIC):

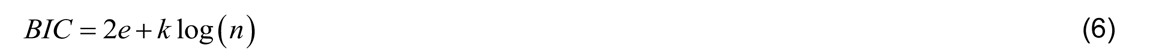

where *k* was the number of free parameters and *n* was the number of trials.

We fit four different model variants including various combinations of the above free parameters. A combination of model validation through parameter recovery analyses and quantitative model comparison converged on a simple 2-parameter normative model fit that included only decision noise (*v*) and subjective hazard rate (^*H*^^) as free parameters (Supplementary Figure 2).

##### Psychophysical kernels

We quantified the time course of the impact of evidence from the main task on choice using logistic regression:

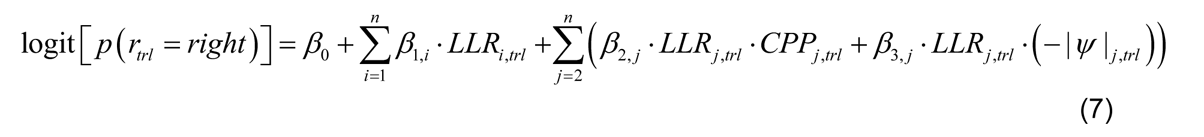

where *i* and *j* indexed sample position within sequences of *n* samples (*n*=10 for this analysis), and *LLR* was the true *LLR*. The dependent variable was participants’ choice (*left*=0, *right*=1) or model choice probability. The set *β*_1_ quantified the impact of evidence at each position on choice and the sets *β*_2_ and *β*_3_ modulations of evidence weighting by *CPP* and -*|ψ|*, respectively. *CPP* was logit-transformed, and both variables were then z-scored. Additionally, all regressors were z-scored across the trial dimension. We fit the logistic regression models using 10-fold cross-validation and an L1 regularization term of *λ* = 0.002, without which convergence issues were encountered for a small number of participants.

We tested for sample positions with above-chance evidence weighting (*β*_1_) and *CPP* (*β*_2_) and -*|ψ|* (*β*_3_) modulations by comparing the group-level distributions of beta weights against zero using cluster-based permutation testing (two-tailed; 10,000 permutations; cluster-forming threshold: *P*<0.01), which corrects for multiple comparisons across sample positions ^83^. For each of the *CPP* and -*|ψ|* modulations, this procedure yielded a large contiguous cluster of sample positions characterized by the expected positive effects (between sample positions 4-10 for *CPP*, and sample positions 3-8 for -*|ψ|*; Figure 2c-e). For analyses of individual differences (across-subject correlations, comparison of sub-groups defined by CAPE scores), we averaged the beta weights within these clusters to compute a scalar modulation score per individual subject; and for such analyses combining *CPP* and -*|ψ|* modulations, we summed across these two scalar values. We also computed a ‘kernel half difference’ measure capturing the relative degree of primacy or recency in average evidence weighting by subtracting the mean of *β*_1,1-5_ from the mean of *β*_1,6-10_. Differences in kernel-based measures across participant subgroups (e.g. lowest and highest quintiles) defined by CAPE P-scores were assessed via two-sample permutation test (two-tailed; 10,000 permutations). Across-participant correlations were assessed via Spearman correlation.

#### Pupillometry

##### Acquisition and preprocessing

Gaze position and pupil diameter from both eyes were recorded at 250 Hz with an SMI RED 500 Eye-Tracker. We restricted our current analysis to pupil data from the main behavioral task. For each task block, we first selected the eye (left *vs* right) giving rise to the pupil signal with the lowest variance across the entire time series and discarded the signal from the other eye from further analysis. Blinks and noise transients were removed from the remaining pupillometric time series using a linear interpolation algorithm in which artifactual epochs were identified via thresholding of the raw pupil size (1.5 mm) and the first derivative of the z-scored time series (threshold=±3.5 zs^-1^). The average time series was then band-pass filtered (0.06-6 Hz, Butterworth), re-sampled to 50 Hz, and z-scored per block. We computed the first derivative of the result, referred to as *pupil* below. Finally, any trial (from onset of the first evidence sample to 5.5 s after trial onset) in which either left or right pupil signals were contaminated by >60% artifactual samples, or any sample forming part of a contiguous artifactual epoch of longer than 1 s, was excluded from all further analysis. For visualization and analysis of the trial-related pupil response in the raw signal (but not it’s derivative), we baseline-corrected the signal for each trial by subtracting the mean pupil size in a 0.1 s window centered on that trial’s onset.

##### Modeling of evoked pupil responses

We assessed the sensitivity of *pupil* to computational variables by segmenting the signal from 0-1 s after sample onset (full-length trials) and fitting:

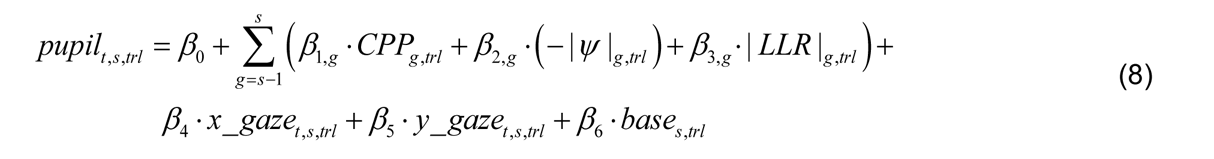

where *t* indicated time point, *x<u>_</u>gaze* and *y_gaze* were horizontal and vertical gaze positions, and *base* was the ‘baseline’ pupil diameter (-0.05-0.05 s around sample onset). |*LLR*| captured a possible relationship with ‘unconditional’ Shannon surprise ^2^. Previous sample *CPP*, -*|ψ|* and |*LLR*| were included because the pupil response is slow, meaning correlations with variables from the previous sample may have caused spurious effects.

We tested for sample-aligned time points with above-chance *CPP* (*β*_1_) and -*|ψ|* (*β*_2_) encoding via cluster-based permutation test (two-tailed; 10,000 permutations; cluster-forming threshold: *P*<0.01). For each of the *CPP* and -*|ψ|* modulations, this yielded a large contiguous cluster of time points characterized by the expected positive effects (between ∼0.4-1.0 s for *CPP*, and ∼0.2-0.45 s for -*|ψ|*; Figure 2c-e). For analyses of individual differences, we averaged the beta weights within these clusters to compute a scalar value capturing encoding strength per individual subject; and for such analyses combining *CPP* and -*|ψ|* modulations, we summed across these two scalar values. Informed by visualization of the trajectories of the average trial-evoked pupil response (Figure 4a), we also computed two scalar measures capturing the magnitude of this response: by averaging the raw (baselined) signal from 0.5-5.5 s following trial onset, and the derivative from 0.2-1.0 s following trial onset. As above for the analysis of kernels, differences in pupil measures across participant subgroups (e.g. lowest and highest quintiles) defined by CAPE P-scores were assessed via two-sample permutation test (two-tailed; 10,000 permutations), and across-participant correlations were assessed via Spearman correlation.

## Data and code availability

Raw behavioral and eye-tracking data will be made available upon publication. Analysis code is available at https://github.com/murphyp7/2024_Murphy_Belief-updating-psychosis-proneness.

## Acknowledgements

We thank Emma Krink, Marlene Petersson, and Sonja Monien for strong engagement in data collection.

This work was funded by the Deutsche Forschungsgemeinschaft (DFG, German Research Foundation) projects DO 1240/4-1, and SFB 936 - Projekt-Nr. A7 (all to THD).

## Author contributions

PRM: Conceptualization, Methodology, Software, Formal analysis, Visualization, Writing – original draft, Writing – review and editing; KK: Conceptualization, Methodology, Software, Writing – review and editing; GM: Investigation, Formal analysis, Writing – review and editing; NK: Investigation; TL: Conceptualization, Methodology, Resources, Writing – review and editing, Supervision; THD: Conceptualization, Methodology, Resources, Writing— original draft, Writing—review and editing, Supervision.

## Supplementary Figures for “Individual Differences in Belief Updating and Phasic Arousal Are Related to Psychosis Proneness”, Murphy et al

**Supplementary Figure 1.**
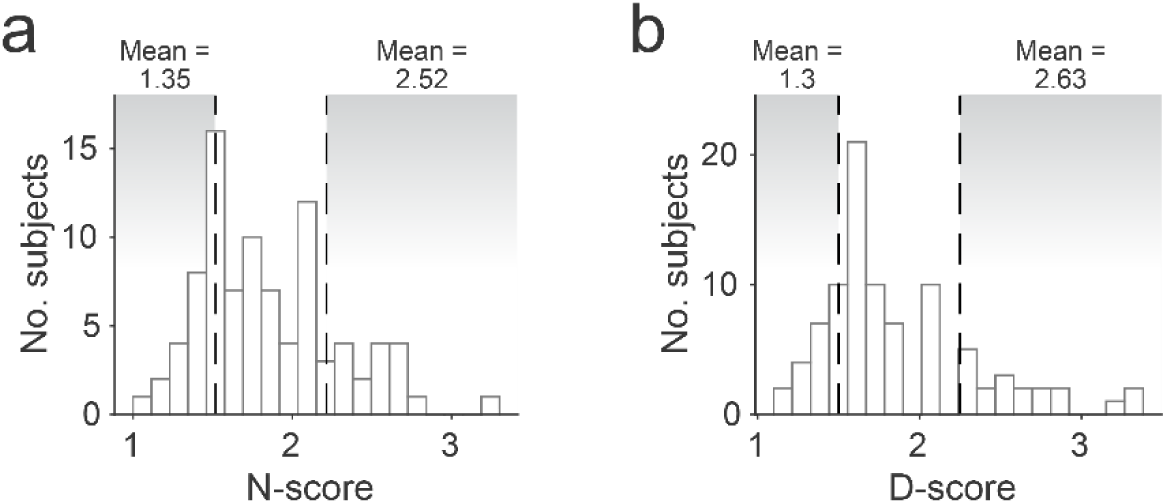
Histograms of N-and D-scores from CAPE in our sample. Like the P-scores shown in main Figure 1e, the N-and D-scores in the community sample also covered a range from healthy to values observed in ultra-high risk samples and in diagnosed patients (Jaya et al., 2021; Mossaheb et al., 2012; Schlier et al., 2015). (a) Distribution of N-scores extracted from CAPE questionnaire data. Vertical dashed lines indicate cutoffs for lowest and highest N-score quintiles; means are mean N-scores within each sub-group. (b) Distribution of D-scores. Format same as in **a**.

**Supplementary Figure 2.**
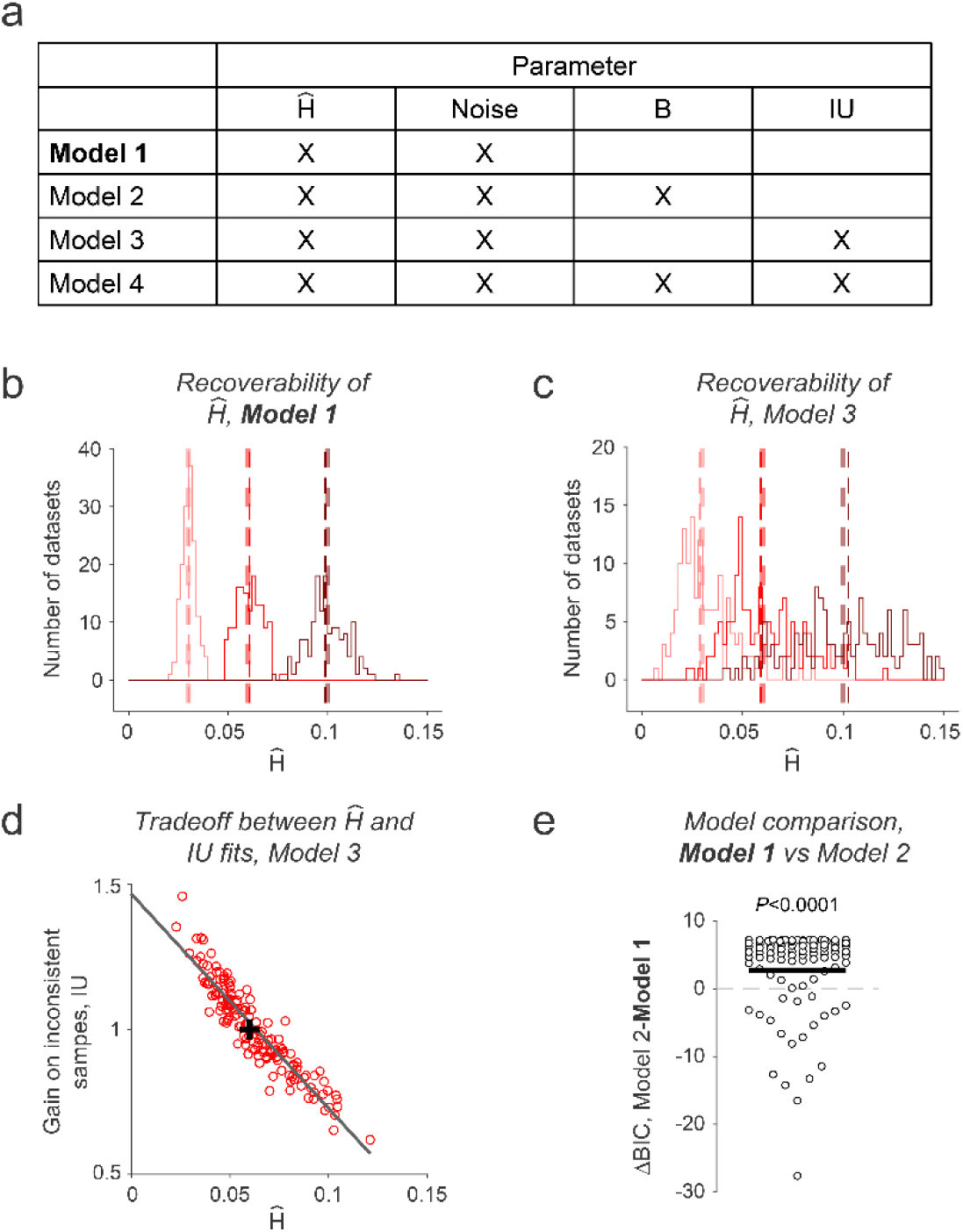
Model validation and comparison favor normative belief updating model with two free parameters, ^H^^ and decision noise. (a) Table specifying free parameters included in each considered model variant. See Methods for descriptions of individual parameters. The model variant reported in the main text, Model 1, is highlighted in bold throughout the figure. (b) Recoverability of the ^H^^ parameter from fits of Model 1. 150 datasets were generated with noise = 2 (close to the group mean value in model fits to our human participants) and ^H^^ set to each of three representative levels (0.03, 0.06 and 0.1; thick vertical dashed lines). Histograms show distributions of recovered ^H^^ parameters (thin vertical dashed lines indicate median ^H^^ per generative parameter set) (c) Same as **b**, but now from fits of Model 3 (which includes a gain on inconsistent evidence parameter, IU, as an additional free parameter). Note the significant increase in the variability of the recovered ^H^^ parameters relative to fits of Model 1 in **b**. (d) Correlation between ^H^^ and IU estimated from fits of Model 3. Cross indicates true generative parameters. Strong negative correlation reflects a tradeoff in fitted ^H^^ and IU parameters, highlighting parameter recovery issue caused by introduction of IU as an additional free parameter. Thus Models 3 and 4 were not considered further. (e) Difference in BIC values for fits of Models 1 and 2 to the participants’ data Positive values indicate support for Model 1, negative values support for Model 2. Although Model 2 provided a better quantitative fit for a modest subset of participants (19 participants with negative values in plot), Model 1 provided the better fit at the group level (P-value, two-tailed permutation test comparing Model 1 and Model 2 BICs).

**Supplementary Figure 3.**
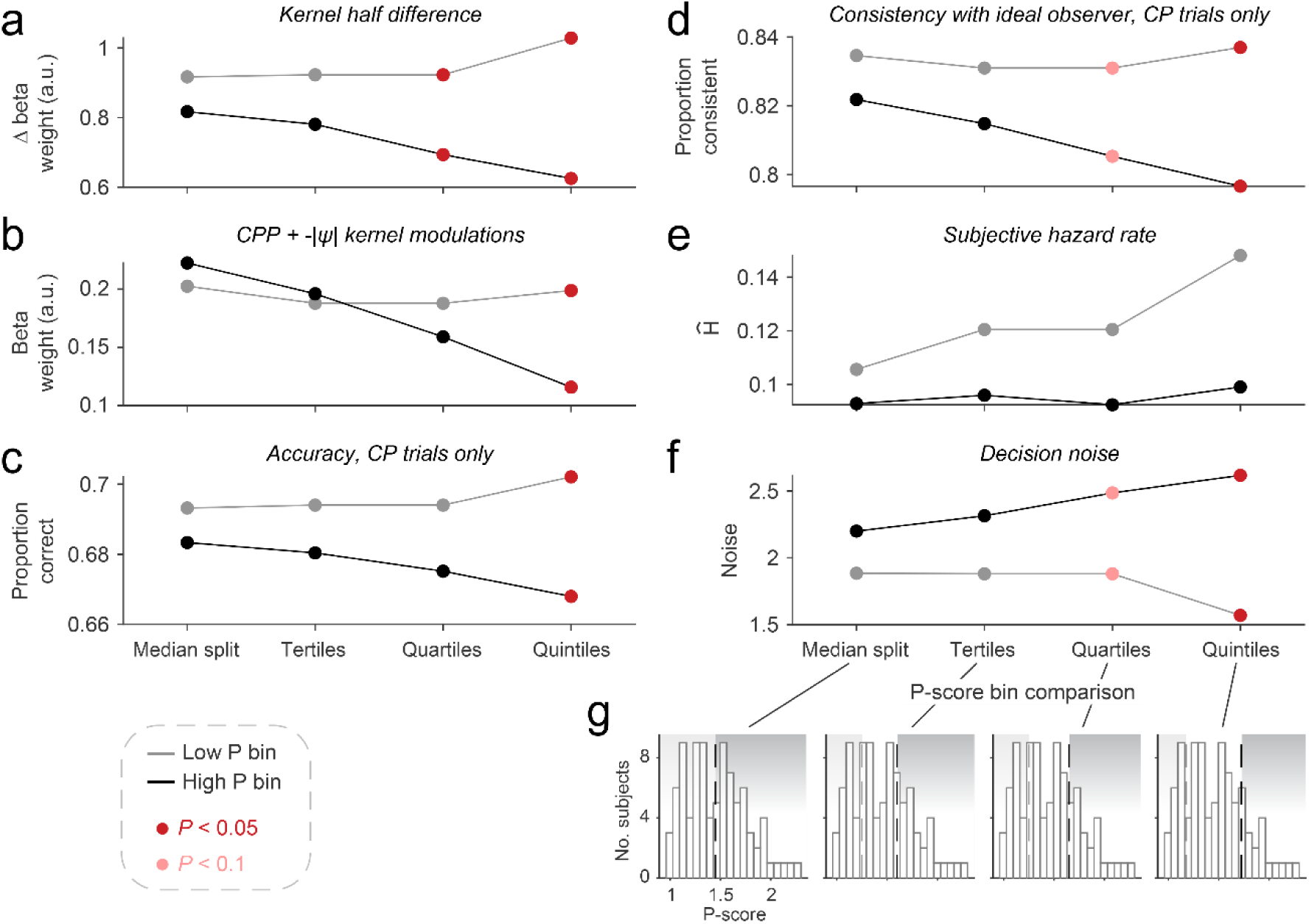
Effect of participant binning on relationships of evidence weighting, behavioral measures and model parameters with individual differences in P-scores. Each plot charts across-participant means of a behavioral or model-based measure, for low (grey) and high (black) P-score participant subgroups defined by each of four different binning procedures (from left to right: median split, first vs. third tertiles, first vs. fourth quartiles, first vs. fifth quintiles). Colored markers highlight statistically significant (dark red, P<0.05) or marginally significant (light red, P<0.1) effects of P-score subgroup for the corresponding binning procedure (two-sample permutation test). (a) Kernel half-difference (subtraction of mean weighting of first 5 samples from mean weighting of last 5 samples) capturing degree of recency in evidence weighting (b) Summed strength of modulations of evidence weighting by CPP (mean modulation weights over sample positions 4-10, significant cluster in Figure 2d) and -|ψ| (mean modulation weights over sample positions 3-8, significant cluster in Figure 2e). (c) Choice accuracy on trials with at least one change-point in task generative state. (d) Consistency of participant choices with those of the ideal observer on trials with at least one change-point in task generative state. (e) Subjective hazard rate parameters from fits of the normative model to participants’ choices (f) Decision noise parameters from fits of the normative model to participants’ choices. (g) Histogram of P-scores from entire sample with cutoffs for low and high P subgroups highlighted for each of the four binning procedures.

**Supplementary Figure 4.**
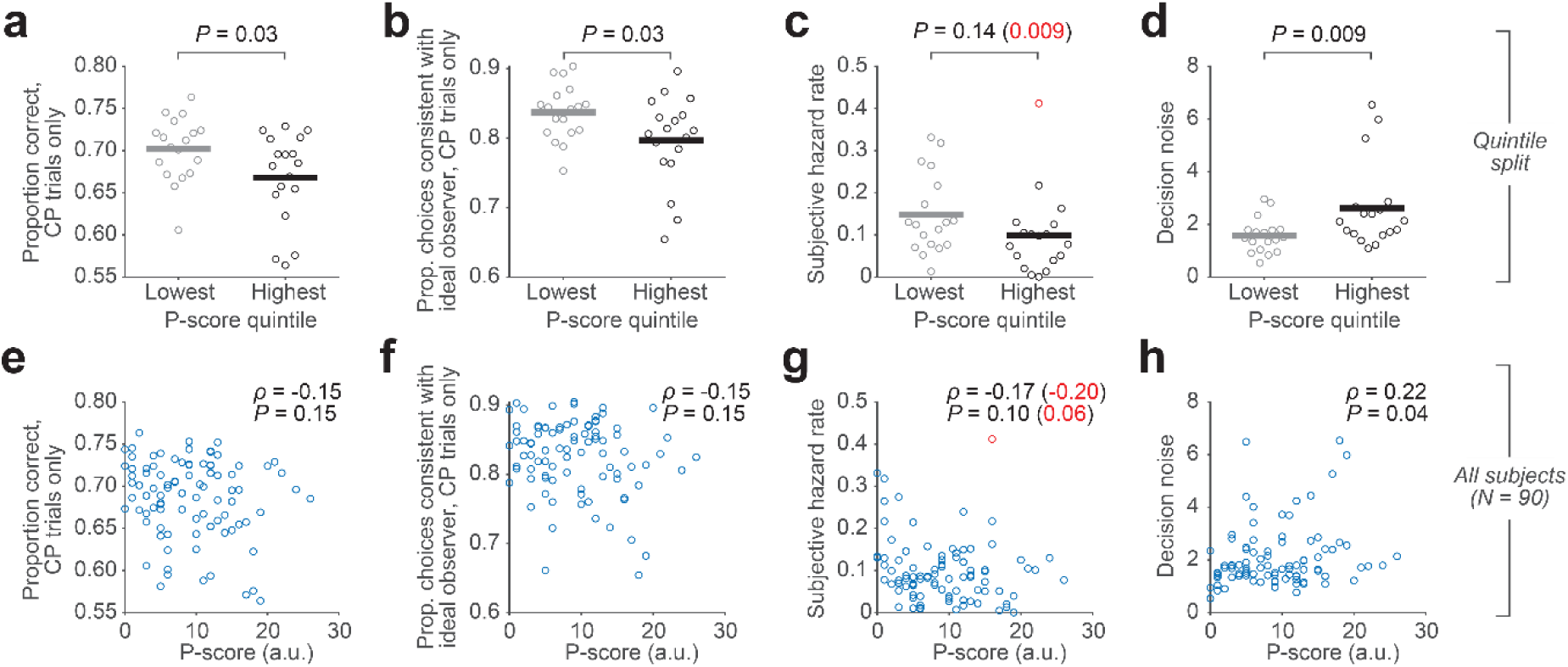
Relationship of overall performance and fitted model parameters with individual differences in P-scores. (a) Choice accuracy on trials with at least one change-point in task generative state, plotted for participants in lowest and highest P-score quintiles. Horizontal lines, mean of data from each subgroup; circles, individual participants. P-value, two-sample permutation test (two-tailed). (b) Consistency of participant choices with those of the ideal observer on trials with at least one change-point in task generative state. Same layout as in **a**. (c) Subjective hazard rate parameters from fits of the normative model to participants’ choices. Same layout as in **a**,**b**. (d) Decision noise parameters from fits of the normative model to participants’ choices. Same layout as in **a**-**c**. (**e-f**) Same performance measures and fitted model parameters as in **a**-**d** but now plotted in scatterplots against P-scores and including all participants (circles). Correlation coefficients and P-values, Spearman correlation. Single participant highlighted in red in **c**,**g** is a potential outlier in the distribution of subjective hazard rate fits. Statistics in parenthesis and red font are from analyses (including re-running of quintile binning, not shown) with this participant excluded.

**Supplementary Figure 5.**
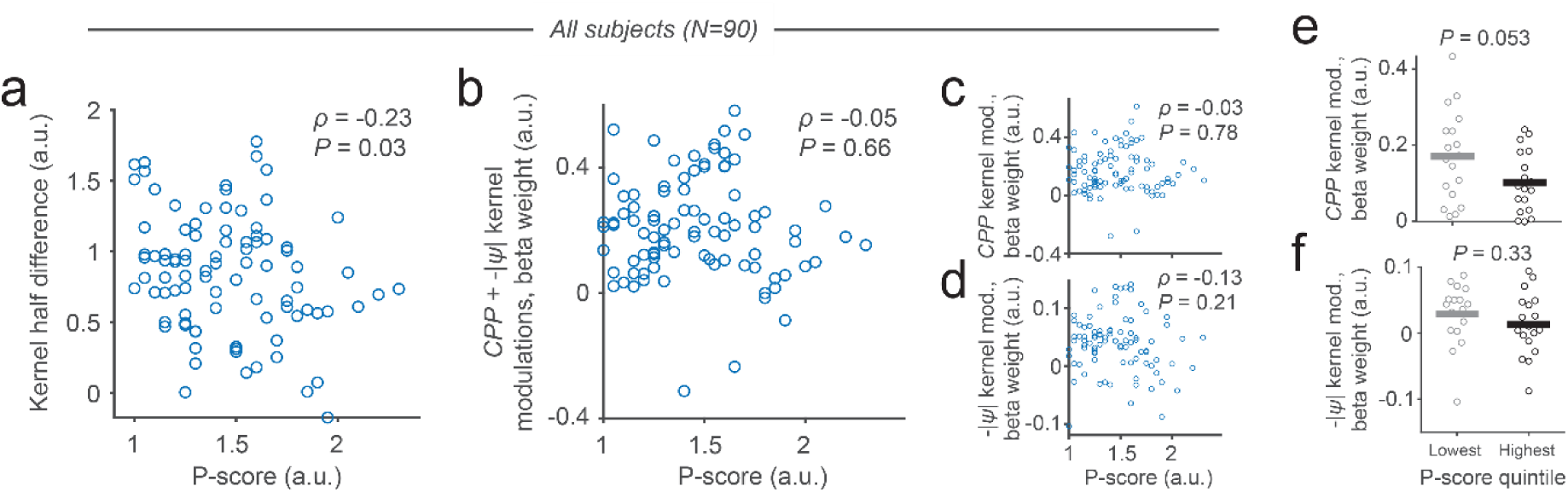
Correlations between kernel metrics and P-scores across all subjects, and decomposition of summed CPP and -|ψ| quintile bin effect. (a) Scatterplot for entire sample (n=90) of P-scores against kernel half-difference for the overall evidence weighting profile. (b) Scatterplot of P-scores against summed strength of modulations of evidence weighting by CPP (mean modulation weights over sample positions 4-10, significant cluster in Figure 2d) and -|ψ| (mean modulation weights over sample positions 3-8, significant cluster in Figure 2e). (**c,d**) Scatterplots of P-scores against individual CPP (**c**) and -|ψ| (**d**) modulations. (**e**,**f**) Individual CPP (**e**) and -|ψ| (**f**) modulations for lowest and highest P-score quintiles. Correlation coefficients and P-values in **a**-**d**, Spearman correlation. P-values in **e**,**f**, two-sample permutation tests (two-tailed).

**Supplementary Figure 6.**
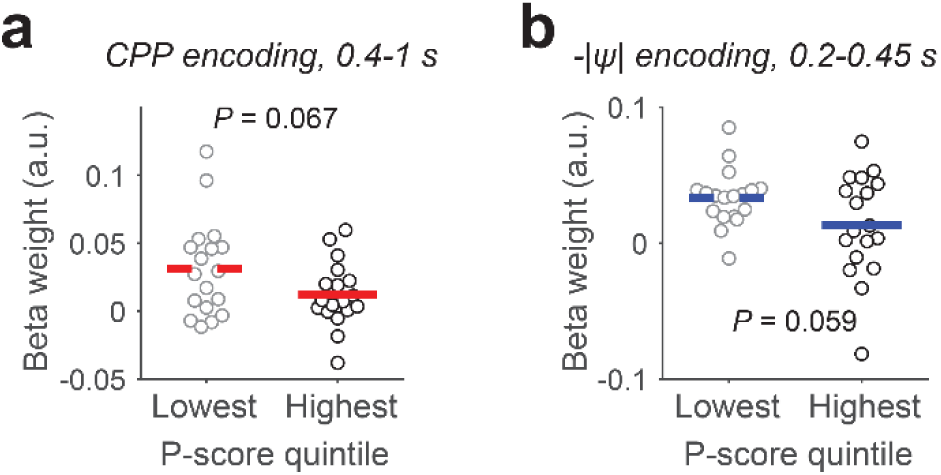
Pupil encoding of change-point probability (CPP) and uncertainty (-|ψ|) in high and low P-score subgroups. (a) CPP encoding. (**b**) Uncertainty encoding. The time windows for each variable were chosen based on the significant encoding in the time-resolved analysis of the whole group (Figure 4d). For both variables, differences between subgroups were marginally significant (permutation tests).

**Supplementary Figure 7.**
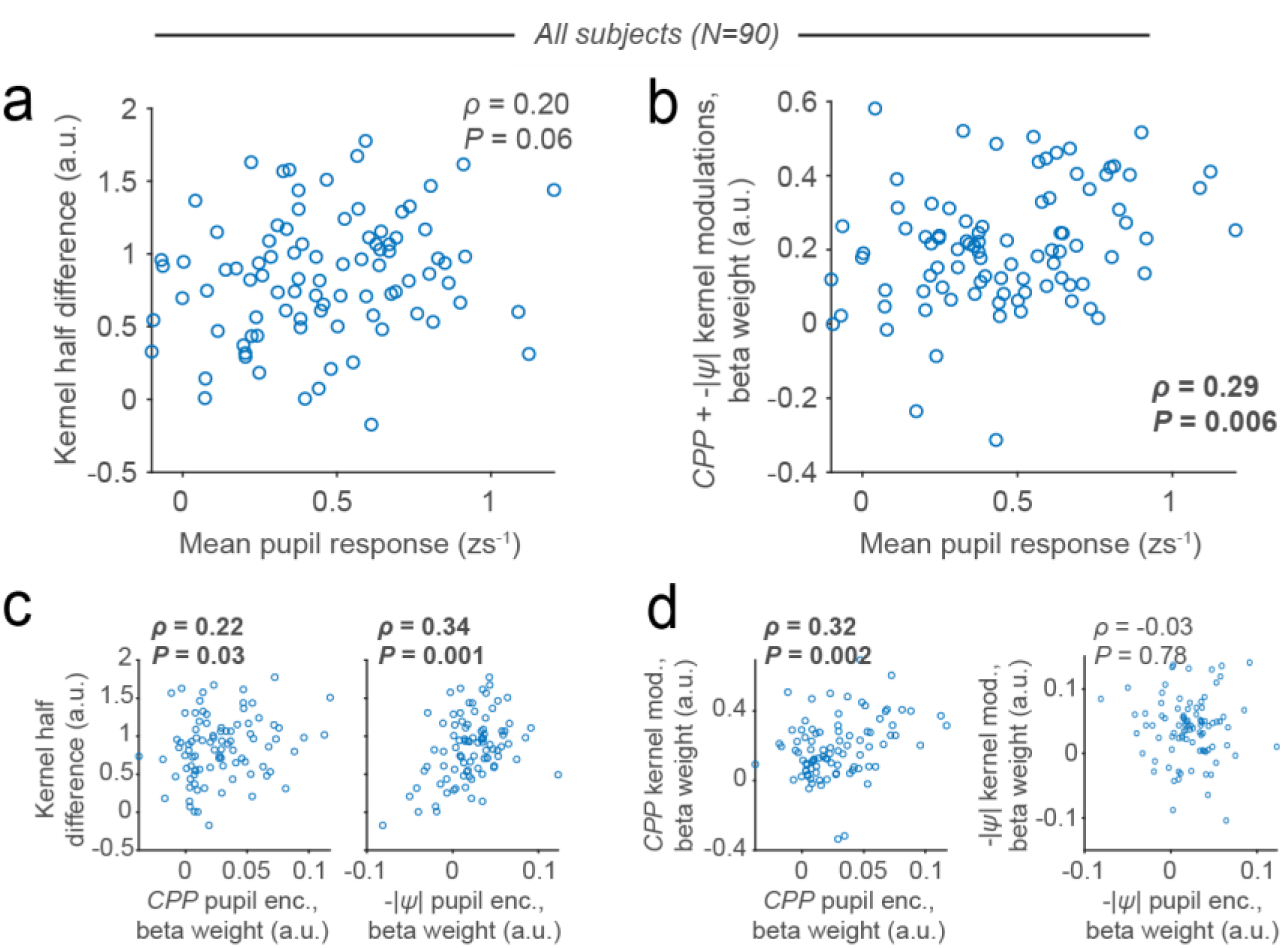
Correlations between kernel and pupil metrics across all subjects. (a) Scatterplot of the mean pupil derivative response (0.2-1 s following trial onset) against the kernel half-difference measure derived from the overall evidence weighting profile. (b) Mean response of the pupil derivative plotted against summed magnitude of CPP (sample positions 4-10) and -|ψ| (positions 3-8) modulations of evidence weighting. (c) Decomposition of correlation reported in Figure 5a, separately plotting kernel half-difference against pupil encoding of both CPP and -|ψ|. (d) Decomposition of correlation reported in Figure 5b, separately plotting CPP modulation of evidence weighting against pupil encoding of CPP, and -|ψ| modulation of evidence weighting against pupil encoding of -|ψ|. Circles, individual participants. Correlation coefficients and P-values, Spearman correlation. Statistically significant (P<0.05) correlations highlighted in bold text.

**Supplementary Figure 8.**
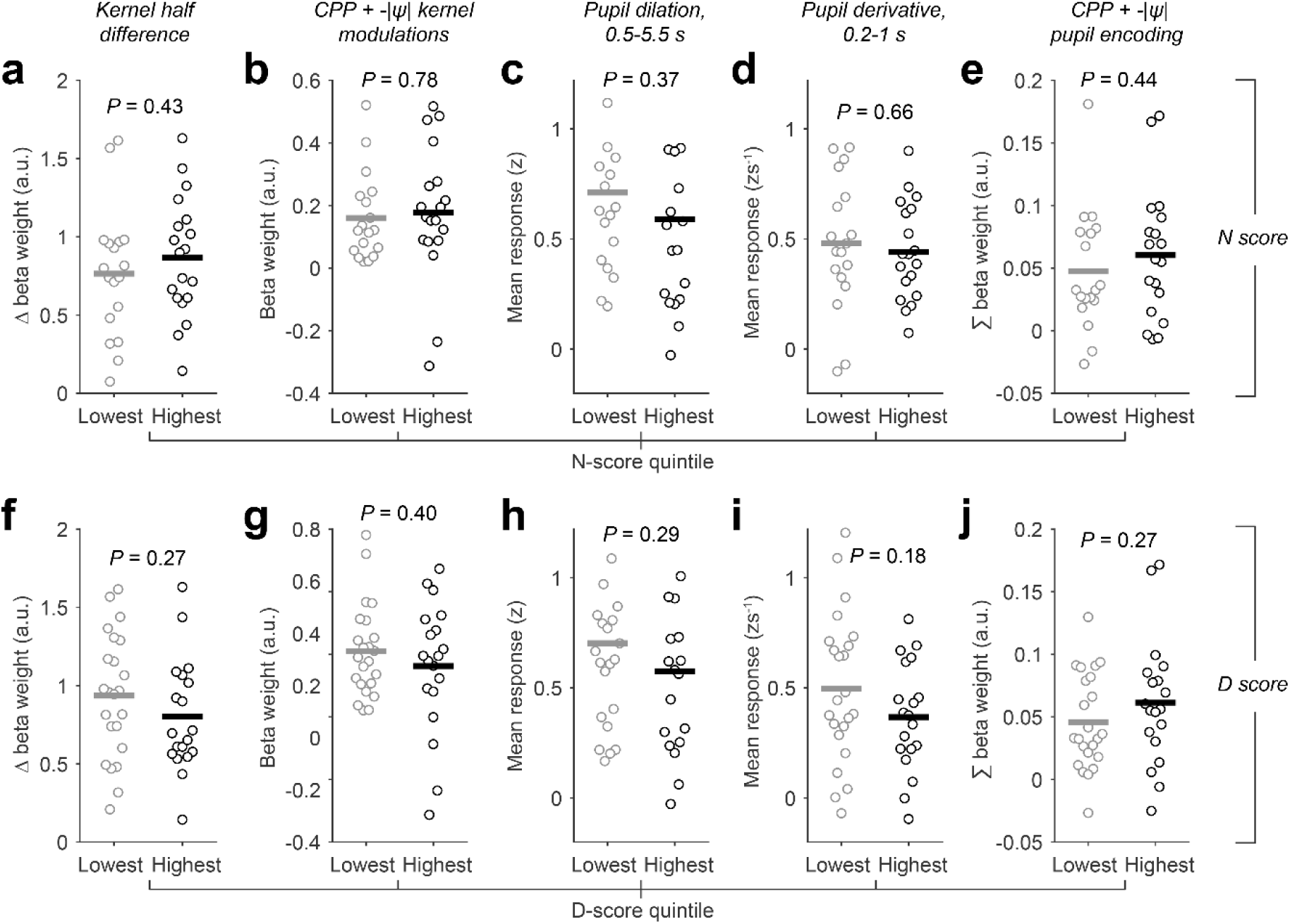
Relationships of evidence weighting and pupil measures to individual differences in N and D scores measured via the CAPE. (a) Kernel half-difference summary measure (subtraction of mean weighting of first 5 samples from mean weighting of last 5 samples) capturing degree of recency in evidence weighting, plotted for participant subgroups defined by lowest and highest N-score quintiles. (b) Summed strength of modulations of evidence weighting by CPP (mean modulation weights over sample positions 4-10, significant cluster in Figure 2d) and -|ψ| (mean modulation weights over sample positions 3-8, significant cluster in Figure 2e), plotted for lowest and highest N-score quintiles. (c) Overall trial-related pupil response for lowest and highest N-score quintiles. (d) Early response of the pupil first derivative for lowest and highest N-score quintiles. (e) Encoding of change-point probability (CPP) and uncertainty (-|ψ|) in pupil responses (pooled) for lowest and highest N-score quintiles. (**f-j**) Same format and dependent variables as **a**-**e**, but now with participant subgroups defined by lowest and highest quintiles of the D-score distribution. Horizontal lines in all panels, mean of data from each subgroup; circles, individual participants. P-values, two-sample permutation tests (two-tailed).

**Supplementary Figure 9.**
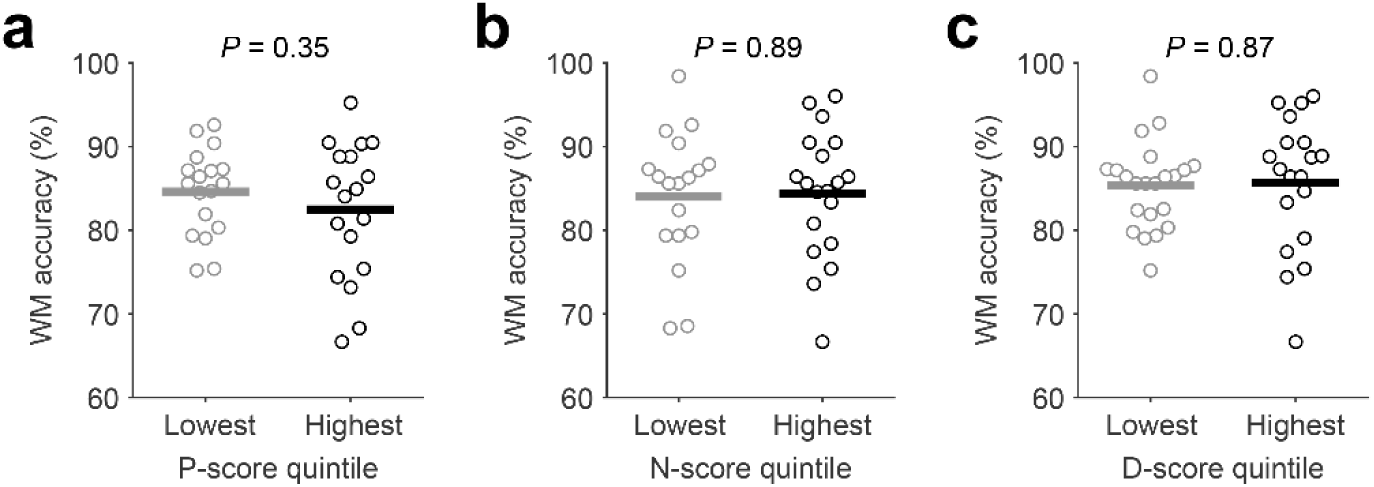
Relationships of CAPE scores to performance on delayed match-to-sample working memory task. (a) Accuracy of ‘same’/’different’ reports on delayed match-to-sample working memory task, plotted for participant subgroups defined by lowest and highest P-score quintiles. See Methods for details on delayed match-to-sample task. (b) Same as **a**, but now with participant subgroups defined by lowest and highest quintiles of the N-score distribution. (c) Same as **a,b**, but now with participant subgroups defined by lowest and highest quintiles of the D-score distribution. Horizontal lines in all panels, mean of data from each subgroup; circles, individual participants. P-values, two-sample permutation tests (two-tailed).

